# Disruptive effect of rainfalls on the diurnal periodicity of airborne wheat rust spore under field conditions

**DOI:** 10.1101/2024.10.03.616546

**Authors:** Frederic Suffert

## Abstract

Stripe rust and leaf rust, caused by *Puccinia striiformis* f. sp. *tritici* (*Pst*) and *Puccinia triticina* (*Pt*), respectively, are major threats to wheat production. Forecasting epidemics requires a deeper understanding of the mechanisms driving spore dispersal. Many studies have either employed field data for purely correlative approaches without incorporating established knowledge on physical mechanisms or, conversely, relied on specific physical approaches in controlled environments focusing on only a few mechanisms or factors. Little emphasis has been placed on holistic field-based studies, where wind and rain play crucial roles. This study fills that gap by attempting to unravel the processes by which rainfall affects airborne spore concentrations over a wheat canopy during active rust epidemics. Over more than two months, bi-hourly spore counts from Burkard traps were integrated with detailed meteorological data, revealing both seasonal and diurnal trends. Diurnal peaks in airborne spore concentrations, typically driven by cyclic changes in wind and humidity, were dramatically altered by rain. Rain events either amplified spore concentrations by up to 25-fold through ‘rain-puff’ and/or depletes them via ‘wash-out’ and ‘wash-off’. Rains events from the dataset were classified into categories with distinct impacts: ‘precursor’ rains often trigger spore release, while ‘follower’ (and prolonged rains) reduce airborne spore concentrations. Moreover, differences in the dispersal dynamics of *Pst* and *Pt* were observed, and some were linked to how humidity and wind influence spore clustering. These results provide valuable insights for a more integrated understanding of the effect of rain and in order to enhance forecasting models.

## Introduction

Wheat rusts are harmful in many wheat-growing regions worldwide. Stripe rust caused by *Puccinia striiformis* f. sp. *tritici* (*Pst*), and, to a lesser extent, leaf rust caused by *Puccinia triticina* (*Pt*), are the most problematic in Europe (Huerta-Espino et al., 2011; Hovmøller et al., 2011). Stem rust (*Puccinia graminis* f. sp. *tritici*) is considered to be re-emerging there (Lewis et al., 2024).

Rust epidemics results from repeated infection cycles that occur within a field canopy from leaf to leaf, driven by the spread of spores liberated by pustules (uredinia). Most of the spores are dispersed by wind and rain over varying distances in short periods of time during a growing season (Aylor, 1990; McCartney and Fitt, 1998). When a spore is deposited on a wheat leaf and phyllosphere conditions are favorable, it gives rise to new uredinia then sporulates and acts as sources of secondary inoculum (Sache et al., 1993). Spores are also dispersed over larger scales, driven by long-distance air mass movements. Such dispersal creates recurrent ‘*Puccinia* paths’, a phenomenon resembling annual migrations, documented for stripe rust in North America (Chen, 2002) and China (Hu et al., 2020), as well as for stem rust in in India (Nagarajan & Singh, 1990), North America (Kolmer 2001), East Africa (Meyer et al., 2017) and, more recently, in Europe (Lewis et al., 2024). Long-distance dispersal can also lead to stochastic founder effects in wheat rust populations, resulting in the introduction of exotic genotypes in new territories (Brown and Hovmøller, 2002).

Since it is almost impossible to mark and track individual spores, their dispersal is usually inferred indirectly by ground-based airborne inoculum monitoring at specific location without knowing their precise origin (Mahafee et al., 2023). Spore traps, such as Burkard or Rotorod, allows to estimate the amount of spores per unit volume of air sampled over a given time period (West and Kimber, 2015; Crisp et al., 2013). This requires coupling the airborne inoculum sampling with diagnostics based on optical or molecular methods (e.g., qPCR). Tracking changes in airborne spore concentration throughout a growing season is useful for feeding into models to forecast epidemic dynamics. However, the relationship between airborne spore concentrations above a wheat canopy and the disease severity assessed a few days or weeks later is not straightforward. High concentrations generally lead to numerous infections, and reciprocally, numerous infections produce large quantities of dispersible spores. However, the impact of a few spores—which might initiate epidemics in distant fields—is less predictable.

Spore dispersal processes involved at short distance during a rust epidemic is traditionally divided into three successive stages: removal from uredinia, transport through the atmosphere, and deposition on leaves (McCartney, 1994; Sache, 2000). Extensive research has been conducted to characterize the physical mechanisms driven each stage, as well as the influence of meteorological variables—primarily wind and rain, but also relative humidity and sun radiation.

Wind is one of the main meteorological variables influencing the removal and transport of rust spores. Experiments conducted in a wind simulator (Fitt et al., 1986), which created conditions close to real-life scenarios, showed that the amount of removed spores was positively correlated with wind velocity, but that low wind velocities (0.5 m.s^-1^) do not hinder short-distance dispersal (Srivastava et al., 1987; Geagea et al., 2000). In contrast, experiments conducted in miniaturized wind tunnels or centrifuges, which are less realistic, revealed threshold wind velocities necessary for spore release, with variations between species: lower for *Pt* (between 0.7 and 1.3 m.s^-1^) (Geagea et al., 1997) and *P. graminis* (between 0.5 and 1.5 m.s^-1^) (Smith, 1966) compared to *Pst* (between 1.3 and 1.8 m.s^-1^) (Geagea et al., 1997). The difference in the forces required to remove spores was related to the size of the dispersal unit, which is larger in *Pst* than in *Pt* due to spore clustering (Rapilly, 1970; Rapilly, 1979). For *Pt* and *Pst*, at high wind velocities exceeding 2.3 m.s^-1^ and 2.8 m.s^-1^ respectively, nearly 90% of the spores were released within the first 10 seconds. At moderate wind velocities, the proportion of removed spores increased with wind duration, from 10 to 60 seconds (Geagea et al., 1997). These results obtained with constant wind velocities should be interpreted with caution as they do not fully account for wind gusts or turbulences. The leaf surfaces in a canopy are represented by agrophysicists within a laminar boundary layer of relatively slow-moving air, which protects spores from being removed and is disrupted by wind gusts (Shaw and McCartney, 1985; Aylor et al., 1990).

Rain, another crucial factor in the removal, transport, and deposition of wheat rust spores, has been also extensively studied. Its impact was long underestimated due to the hydrophobic nature of the spores. Early field investigations have suggested that rainfalls have more complex effects on spore movement than wind (Hirst, 1961; Rapilly, 1979). Specific experimental devices, such as a rain tower (Fitt et al., 1986), have shown that when a raindrop strikes a uredinium on a leaf, spores are incorporated into the splash droplets and dispersed beyond the impact zone (Huber et al., 1996). The efficiency of this short-distance dispersion, known as ‘rain-splash’, is an increasing function of the kinetic energy of the raindrops, and consequently depending on the intensity and duration of the rain (Geagea et al., 1997). Additionally, when a drop impacts, it induces the dry dispersion of spores from uredinia near the point of impact through mechanical energy transfer, a process known as a ‘rain-puff’ (Hirst and Stedman, 1963; Rapilly et al., 1970). By generating a range of raindrop diameters corresponding to real rainfalls (2.5-4.9 mm) and falling from heights of 5 to 100 cm, Geagea et al. (1999; 2000) showed that more spores of *Pst* and *Pt* were rain-splashed than dry-dispersed. Dispersal efficiency dropped to nearly zero after a 10-minute rainfall with an intensity of 40 mm.h^-1^ for both rusts. This was due to the exhaustion of uredina and the fact that restoration of sporulation capacity requires a dry period, estimated of 2 hours for *Pt* and up to 6 hours for *Pst* (Geagea et al., 1999), with the difference between the two rusts once again attributed to the spore clustering of the latter (Rapilly et al., 1970). These results, complemented by physical modelling studies (Huber et al., 1996; Saint-Jean et al., 2004), allow for qualitative predictions of the dispersal ability of different types of rain. Very small drops, due to their reduced kinetic energy, contribute minimally to spore removal compared to the relatively rare large drops with high kinetic energy. Experiments conducted by Kim et al. (2019) on *Pt* demonstrated that dry dispersal away from the wheat leaf boundary layer is also possible through raindrop-induced vortex flows. Wu et al. (2024) revealed that spore removal can result from vibrations induced by raindrops, which generate a lateral flow stream upon impacting leaves that are not rigid but rather flexible structures (Gart et al., 2015). Jumping-droplet condensation is another passive mechanism that has recently been shown to propel *Pt* spores up to 5 mm from superhydrophobic wheat leaves (Nath et al., 2019). This dispersal is further enhanced by a gentle breeze in the absence of rain or strong winds (Mukherjee et al., 2021).

The role of rain in spore deposition on wheat leaves through ‘wash-out’ was suggested by the presence of rust spores in rainwater (Rowell and Romig, 1966; Nagarajan et al., 1977). This mechanism was quantified by collecting rainwater during 10-minute intervals in glass vials placed a few meters from wheat plots. The waterborne spore concentration, ranging from 300 to 30.000 spores.cm^-3^, increased with airborne spore concentration and decreased with the rain duration (Suffert, 1999; Sache et al., 2000). To add another layer of complexity, ‘wash-off’ of rust spores already deposited on leaves by incident raindrops was demonstrated placing wheat seedlings either sheltered from or exposed to rainfalls of varying durations and intensities. This resulted in significant differences in *Pt* infection during a violent thunderstorm (60 min) and a shower (10 min), but minor differences with lighter rains (Suffert, 1999; Sache et al., 2000). While the effects of rain can be beneficial for spore removal and deposition, they can ultimately be detrimental to rust infection, as also exemplified in coffee rust caused by *Hemileia vastatrix* (Avelino et al., 2020).

Correlative approaches have been employed in field conditions to evaluate the combined effects of climate conditions on the temporal patterns of airborne spores for various fungal taxa, including cereal rusts (e.g., Grinn-Gofroń et al., 2020). Although most studies have offered valuable insights, some conclusions have been artificially derived due to reasoning challenges and difficulties in disentangling complex interrelationships between variables. For instance, the significant influence of average air temperature can be overstated when the time scales used do not align with the previously reviewed physical mechanisms. This was observed in a study analyzing the relationship between daily airborne spore concentrations up to three days prior and the corresponding day’s meteorological variables (Sánchez Espinosa et al., 2023), as well as in another study examining the relationship between daily airborne spore concentrations collected over a longer period (7 days) and average daily temperature (Hu et al., 2024). Such approaches, while stating an obvious correlation between spore release, epidemic dynamics, seasonal wheat growth cycle, and rising seasonal temperatures, may obscure the more complex impact of other variables that better explain the underlying physical processes. The framework proposed by Chamecki et al. (2012) is highly useful in overcoming this bias. However, the time intervals employed—in their case, extremely short (spore trapping every 30 seconds for 30 minutes)—can present a practical limitation due to the scarcity of data at this time step.

The current study collected high-resolution data on airborne spore concentrations of *Pt* and *Pst*, as well as meteorological data, to characterize how key climate factors—wind, humidity, and especially rain—affect spore dispersion mechanisms above a wheat canopy. The goal was to understand the complexity of their effects in light of the previously reviewed physical mechanisms.

## Material and methods

### Experimental device

The experimental setup consisted of two rectangular wheat plots (50 × 100 m), located on the INRAE experimental domain in Grignon (Yvelines, France). The plots were situated on a windy, open plateau, highly exposed to both precipitation and sunlight. The plots were sown in the autumn 1998 with two varieties of wheat: ‘Victo’, highly susceptible to stripe rust but resistant to leaf rust, and ‘Soissons’, susceptible only to leaf rust. To initiate the development of each rust epidemics, two contaminating pots of seedlings infected with *Pst* (French pathotype 6E16 virulent on *Yr2, Yr6, Yr7*, and *Yr8*; de Vallavieille-Pope et al., 2012) were placed into the first plot on April 1st, 1999, and twenty pots of seedlings infected with *Pt* (isolate B9384-1C1 virulent on *Lr14a* and *Lr13*) were placed into the second one.

### Monitoring of epidemic development

The development of both epidemics wad evaluated through weekly measurements of disease severity, conducted from April 27 to June 10, 1999 for stripe rust, and from May 18 to June 21 for leaf rust. For the sake of representation, the dates were subsequently converted to Day of Year (DOY). The disease severity was assessed by estimating the leaf area occupied by the uredinia (**Figure 1**) on the top three leaf layers of five steams taken at ten representative points within each plot. These estimates were made visually with the support of the ‘Distrain’ software (Tomerlin and Howell, 1988).

**Figure 1.**
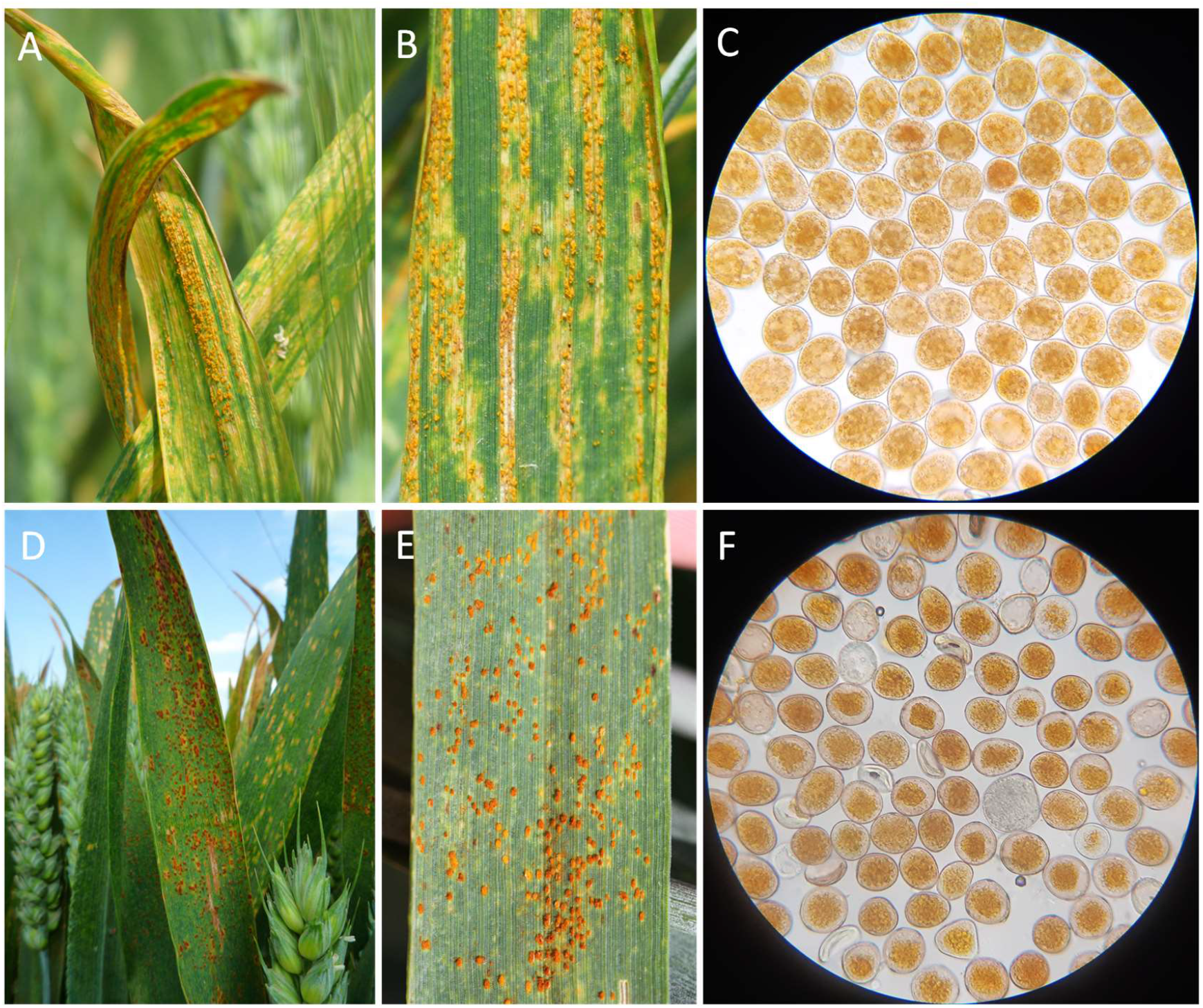
Wheat leaves infected by **(A)** *Puccinia striiformis* f. sp. *tritici* (*Pst*, stripe rust) and **(D)** *Puccinia triticina* (*Pt*, leaf rust), displaying **(B)** stripes of uredinia and **(E)** partially clustered uredinia, respectively. The urediniospores can be distinguished under the microscope by their color and size: **(C)** yellow-orange, measuring 27±2 × 23±2 μm for *Pst*; **(F)** orange-brown, measuring 24±3 × 20±2 μm for *Pt* (400x magnification).

### Spore trapping and estimation of airborne spore concentration

The airborne concentration of *Pst* and *Pt* spores was estimated using two volumetric Burkard spore traps (Burkard Manufacturing Co Ltd, Rickmansworth, UK) installed in each plot, located 25 meters from where the contaminating pots were placed. From April 22 to July 10, 1999, spores were trapped on a rotating drum covered with a transparent Melinex polyester tape (14 mm wide x 340 mm long) coated with a blend of petroleum jelly and wax. The drum completed a full rotation every 7 days at a speed of 2 mm.h^-1^, with an air intake rate of 0.6 m^3^.h^-1^ (or 10 L.min^-1^) maintained throughout the duration of the trapping (Gregory, 1954). Temporal calibration was achieved by depositing Lycopodium powder in front of the sampler air intake after each weekly replacement of the drum, marking the start of each new tape. It was assumed that all spores contained within 1.2 m^3^ of air were captured over a 2-hour interval and adhered to a 4 mm wide section of the tape. Each week, the tape was collected and divided into seven 48 mm long sections, one for each day, using a graduated ruler. These sections were mounted between a microscope slide and cover slip using a Gelvatol solution. In each daily section, spores were identified and counted under a light microscope. The spores of each species were distinguished by their color and size (**Figure 2**). For each section, 12 vertical traverses, each 355 μm wide (corresponding to the microscope’s field of view at 200x magnification) and spaced 4 mm apart, were visually scanned. These traverses were considered representative of the spores trapped during each 2-hour interval (**Figure 3**).

**Figure 2.**
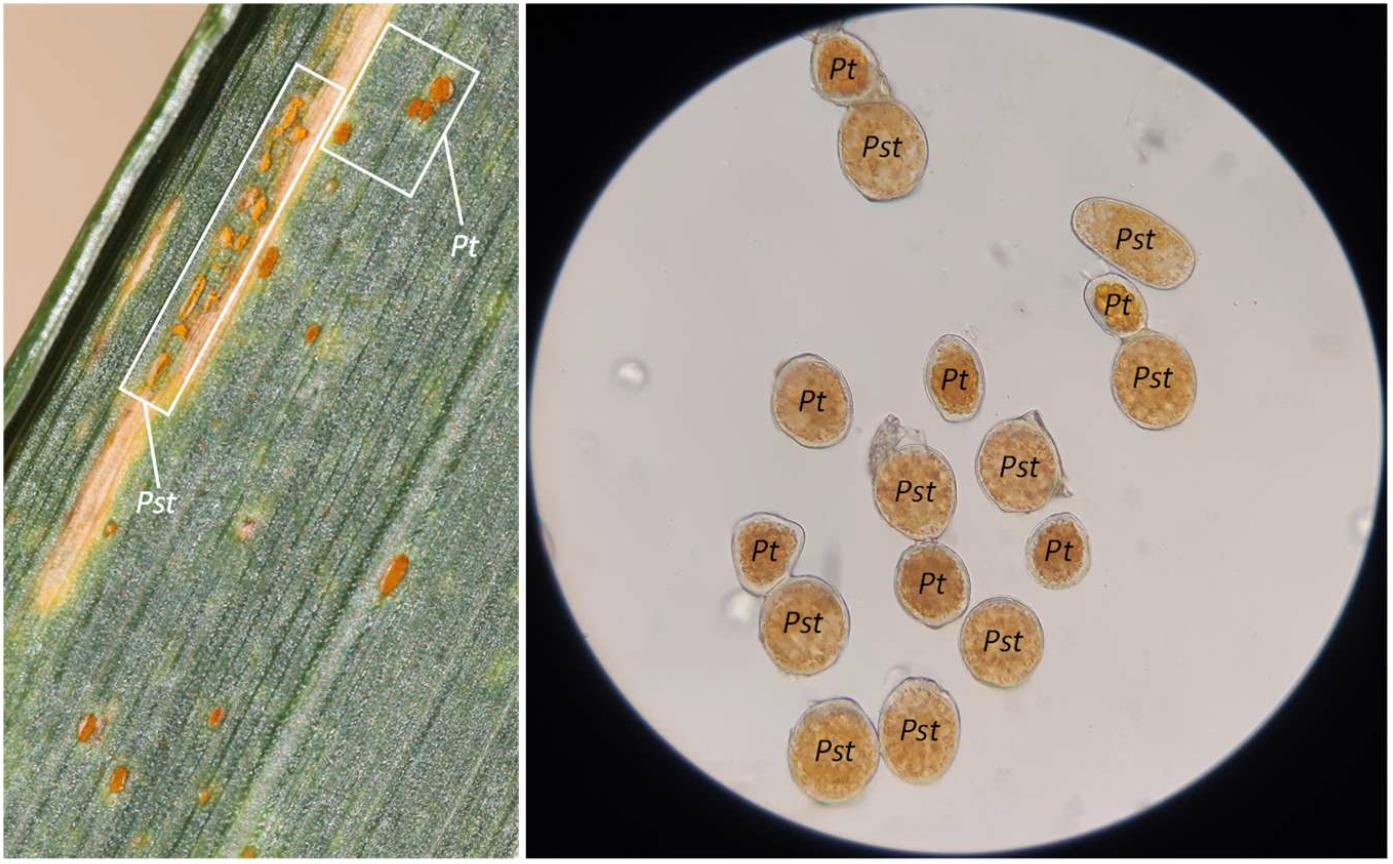
**(A)** Co-occurrence of uredinia from both stripe rust (*Puccinia striiformis* f. sp. *tritici, Pst*) and leaf rust (*Puccinia triticina, Pt*) on the same wheat leaf and **(B)** urediniospores collected on a tape from a Burkard spore trap, observed under a microscope (400x magnification), distinguishable by their size and color.

**Figure 3.**
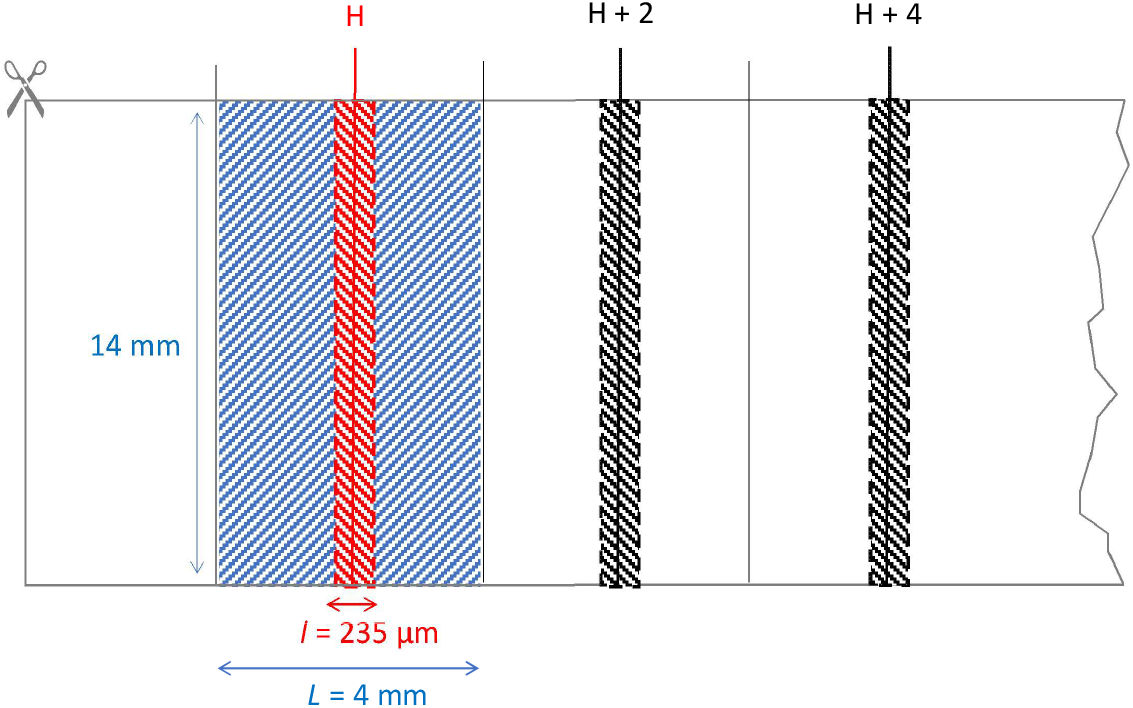
A section of tape from a Burkard trap showing the area where spores were trapped during the 2-hour interval centered on hour H (blue), and the specific traverse used for spore counting (red).

The airborne spore concentration (*C*) was estimated (flux, in m^3^.2h^-1^) à using the following formula:

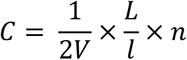

where *V* is the volume of air sampled (0.6 m^3^.h^-1^), *L* is the width of the tape section where the spores sampled during the 2-hour interval adhered (4 mm), *l* is the width of the traverse (235 μm), and *n* is the number of spores counted within the traverse.

In total, *Pst* spores were counted on 556 traverses from April 22 (DOY 112) to June 7 (DOY 158), while *Pt* spores were counted on 546 traverses from May 26 (DOY 146) to July 10 (DOY 191), both corresponding to 46 days of trapping. Due to the staggered timing of the two rust epidemics, the spores of both species were present on the same tape only between DOY 146 and DOY 158, which minimized the risk of confusion and facilitated counting throughout the trapping interval.

### Meteorological data

Meteorological data were continuously collected from April 19 to July 10, 1999 using an automated Campbell CR 10 micrometeorological station (Campbell Scientific Ltd, UK). This station was custom-designed at the INRAE Bioclimatology Station and installed in the middle of the ‘Soissons’ wheat plot. Key variables measured at 2 meters above the crop canopy included air temperature (T, in °C), relative humidity (RH, in %), wind velocity (WV, in m.s^-1^) and gust intensity (GI, 10 times the standard deviation of wind velocity measured every 5 seconds over a 15-minute period), and rainfall (R, in mm). The variables were recorded every 15 minutes and then averaged or summed over 2-hour intervals to align with the time step granularity of airborne spore concentration estimation. An Ordinary Least Squares (OLS) regression based on the following model was used to comprehensively quantify the effect of independent meteorological variables on airborne spore concentrations (C):

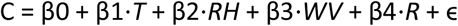

where *T* is the air temperature, *RH* the relative humidity, *WV* the wind velocity, and *R* the rainfall.

Temperature, being a variable traditionally considered in this type of analysis, was included in the model despite its lack of demonstrated impact on spore removal processes (see review in the Introduction) and the potential overlap of its diurnal periodicity with those of RH and WV. GI, being dependent on WV, was excluded from the model.

### Caracterization of rain effects

A ‘rain event’ was defined as a rainfall occurring within a 2-hour interval, regardless of how the rain is distributed within this interval. Each rain event was categorized as either a ‘precursor’ or a ‘follower’. It was considered a precursor if it occurs at least 6 hours after a previous rain event (meaning at least four 2-hour intervals separate the two rain events); otherwise, it was categorized as a follower. Therefore, a prolonged rain interval lasting more than 2 hours may be broken down into an initial precursor event followed by one or several successive follower events (see **Figure 3** for examples). What led to this choice is that the impact of raindrops on airborne spore concentration is known to gradually decrease after the end of a rainfall, which Geagea et al. (1999) estimated to be between 2 hours (the duration defined as a rain event) and 6 hours.

The impact of each rain event on the airborne spore concentration was estimated by calculating the difference between the number of spores trapped during the 2-hour interval of the rain event and the number trapped during the preceding 2-hour interval. Additionally, the difference between the number of spores trapped during the 2-hour interval of the rain event and the following 2-hour interval was calculated. These differences were also calculated for all non-rain intervals (‘No rain’) to estimate the changes in airborne spore concentration that could occur in the absence of rain, serving as a reference value. All differences were normalized, i.e. expressed as a percentage increase or decrease in spore concentration. The first difference provides insight into the ‘suspension effect’ assessed during the rain event, while the second difference provides insight into its ‘wash-out effect’ assessed just after the same rain event. By design, when two rain events occur in succession, the suspension effect of the second rain event overlaps with the wash-out effect of the first one, making it difficult to distinguish which rain event was actually responsible for the change in airborne spore concentration. This change in airborne spore concentration was therefore seen as the result of the action of the beginning of a rain interval (precursor rain) and the continuation of this interval (follower rain). The principle of measuring the impact of each type of rain event on airborne spore concentration, specifically regarding suspension in the air and wash-out dynamics, is illustrated in **Figure 4**. The data processing was carried out for each rust species, *Pst* and *Pt*, during their respective active epidemic phases, defined as the interval from the first day when the total daily number of airborne spores t exceeded 10000 to the last day when this quantity was trapped (**Supplementary Figure 1**).

**Figure 4.**
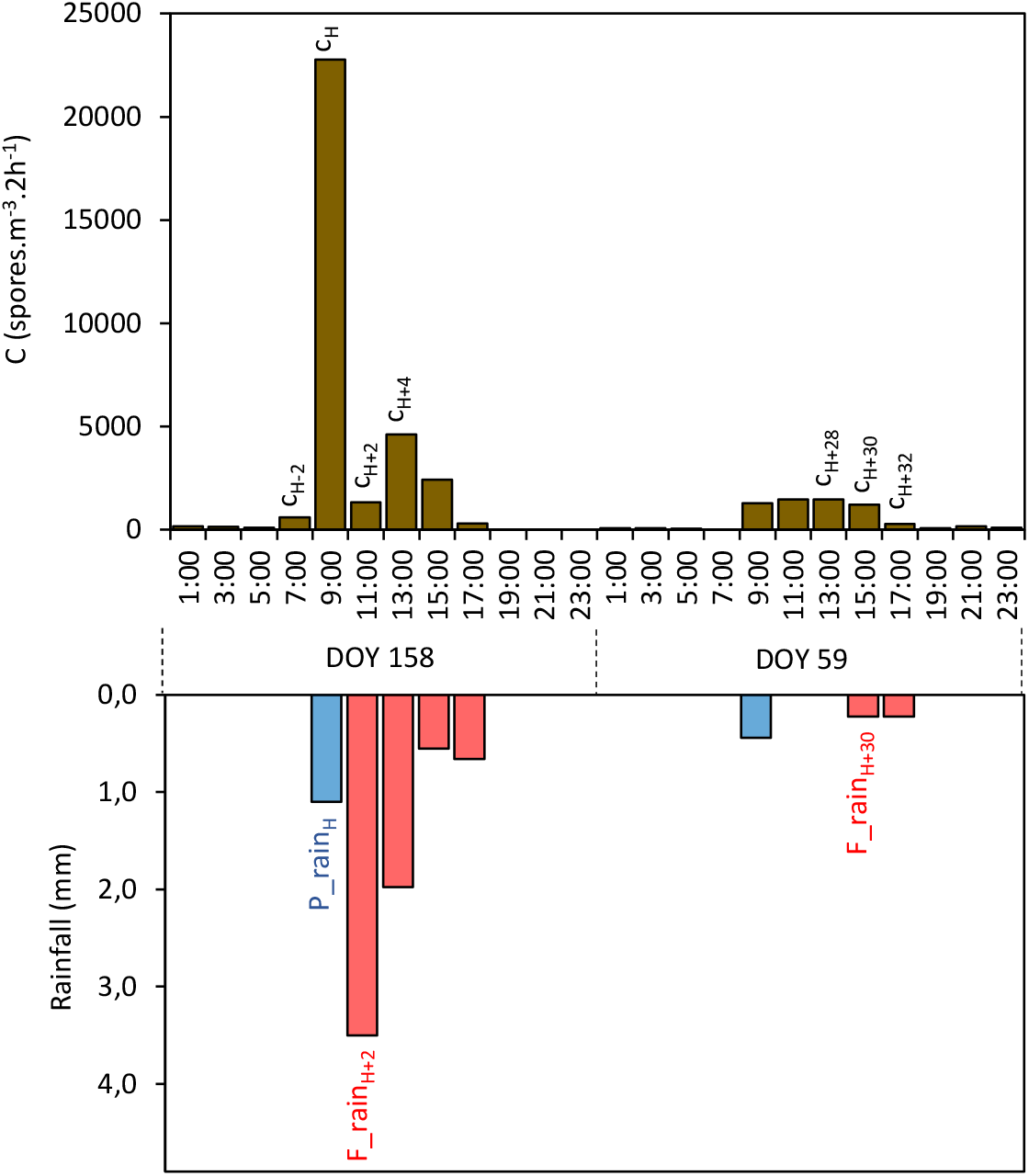
Method for estimating the impact of rain events on changes in airborne *Puccinia triticina* (*Pt*, leaf rust) spore concentrations by distinguishing between ‘suspension effect’ and ‘wash-out effect’ based on the data set presented in **Supplementary Figure 1**. Three case of calculation are here provided for the ‘precursor’ (P_rain_H_) and ‘following’ rain events (F_rain_H+2_) centered on hours H and H+2 on DOY 158 (June 7, 1999), and for the following rain event (F_rain_H+30_) centered on hour H+30 on DOY 159 (June 8, 1999). The upper brown bars represent the cumulative airborne spore concentration (C) during 2-hour intervals. The lower bars represent the amount of water collected during the same intervals, with the blue bars denoting precursor rain—defined as rainfall occurring at least 6 hours after a previous rain event—and red bars denoting following rain, which occurs less than 6 hours after a previous rain event. For P_rain_H_, suspension effect = 100 x (c_H_-c_H-2_)/c_H-2_ and wash-out effect = 100 x (c_H+2_-c_H_)/c_H_. For F_rain_H+2_, suspension effect = 100 x (c_H+2_-c_H_)/c_H_ (thus blending with the wash-out effect of P_rain_H_) and wash-out effect = 100 x (c_H+4_-c_H+2_)/c_H+2_. For F_rain_H+30_ suspension effect = (c_H+30_-c_H+28_)/c_H+28_ and wash-out effect = 100 x (c_H+32_-c_H+30_)/c_H+30_.

### Characterization of humidity and wind effects

The effects of humidity and wind were estimated by analyzing the relationship between the average airborne spore concentration of each rust species and three microclimatic variables (RH, WV, and gust intensity) during the active epidemic phase. Furthermore, the impact of RH on the average size of the dissemination units was tested for each rust species. To this end, the size of each dissemination unit— number of clustering spores, i.e. that overlap or touch each other—was estimated during all 2-hour intervals in which the concentration exceeded 50 spores.m^-3^.2h^-1^. The distribution of dissemination unit sizes was finally retained for analysis in 104 2-hour intervals from JOY 126 (May 6) to JOY 143 (May 23) for *Pst*, and for 96 2-hour intervals from JOY 151 (May 31) to JOY 172 (June 21) for *Pt*.

## Results

### Disease dynamics

Four weeks after the inoculation of *Pst*, on April 27 (DOY 117), several secondary foci were detected in the ‘Victo’ plot. The identification of pathotype 233E137, naturally present in the Île-de-France region in 1999, as well as pathotype 6E16, corresponding to the inoculated isolate also commonly found in the South-West at that time (de Valavieille et al., 2012), suggests that the stripe rust epidemic developed from both artificial inoculation and natural inoculum. It rapidly progressed from these secondary foci, with the entire plot affected by DOY 124. The first leaf rust symptoms displayed on June 1st in the ‘Soissons’ plot with a development of the epidemic from the *Pt* inoculation point before spreading to the rest of the plot. The development of both epidemics was visualized using a logistic model fitted to the average values (power law; Gregory, 1968), which revealed a classic rate of disease progression (**Figure 5A**). The inflection point reached when the airborne spore concentration exceeded approximately 40000 sp.m^-3^.day^-1^ for both rust species (**Figure 5B**).

**Figure 5.**
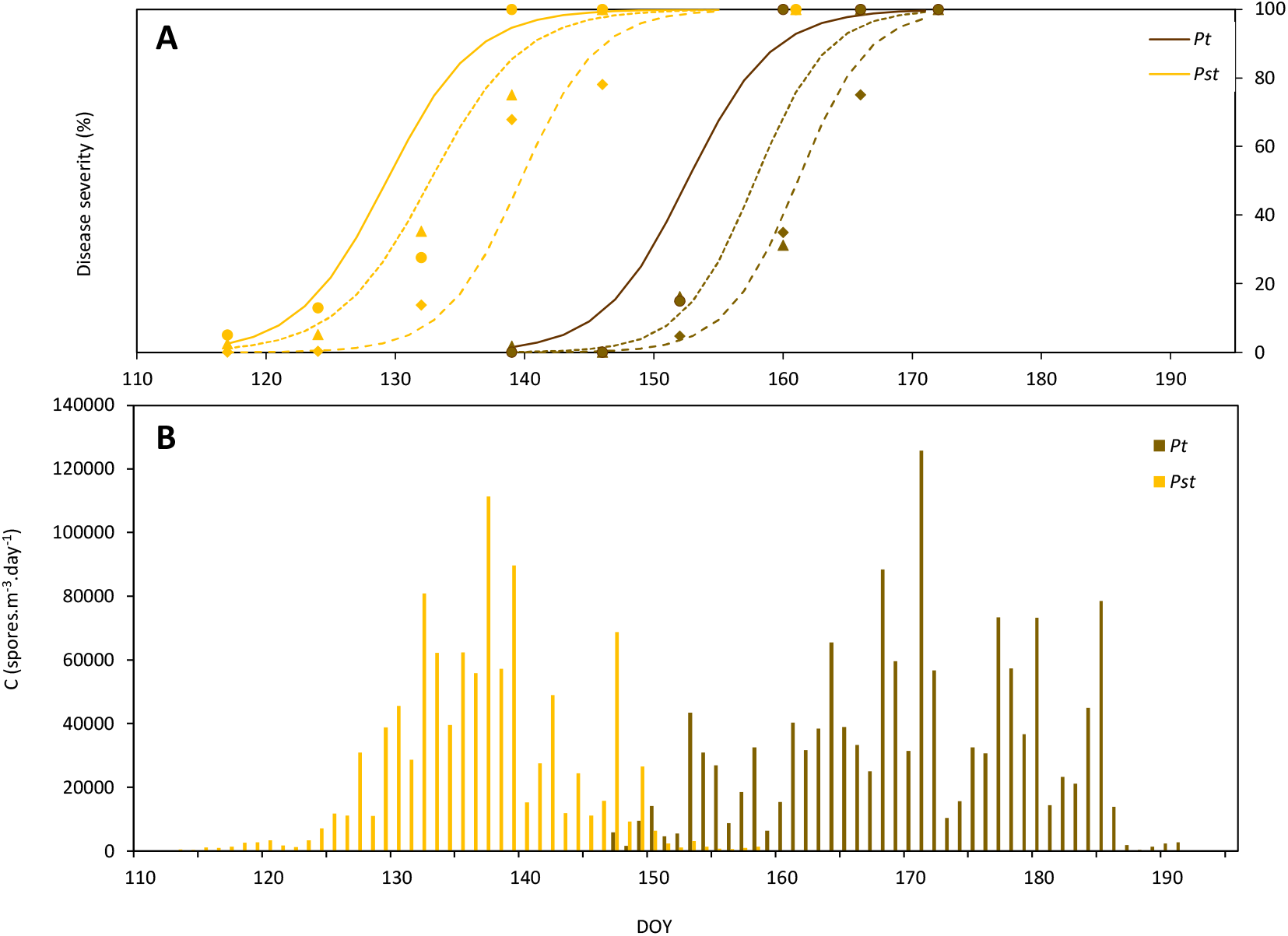
Characteristics of rusts epidemic development in the two wheat plots (Grignon, Yvelines). **(A)** Disease severity (in % of the leaf covered by uredinia) caused by *Puccinia striiformis* f. sp. *tritici* (*Pst*, stripe rust) and *Puccinia triticina* (*Pt*, leaf rust). The points represent the experimental values (average severity on leaves 1, 2, and 3, starting from the top) and the lines depict the fitted logistic curve. **(B)** Cumulative daily airborne spore concentration (in spores.m^-3^.day^-1^) from DOY 115 (April 25, 1999) to DOY 191 (July 10, 1999).

### Airborne spore concentration dynamic

Airborne spore concentration varied greatly from day to day, but a similar overall trend emerged for *Pt* and *Pst*, with a steady increase up to a maximum followed by a steady decline (**Figure 5B**). For *Pst*, the maximum bi-hourly spore concentration was reached on May 19 (YOD 139) at 11 a.m. (> 42000 sp.m^-3^.2h^-1^) and the maximum daily concentration on May 17 (YOD 137) (> 111000 sp.m^-3^.day^-1^) (**Supplementary Figure S1**). For *Pt*, the maximum bi-hourly concentration was reached on June 17 (YOD 168) at 11 a.m. (> 32000 sp.m^-3^.2h^-1^) and the maximum daily concentration on June 20 (YOD 171) (> 125000 sp.m^-3^.day^-1^).

An analysis of intra-day airborne spore concentrations was conducted using data from the active epidemic phase. A diurnal periodicity in the spore dispersal was identified between May 5 (YOD 125) and May 29 (YOD 149) for stripe rust (**Figure 6A**), and between May 30 (YOD 150) and July 5 (YOD 186) for leaf rust (**Figure 6B**). During both epidemics, nearly 80% of the spores were released between 8 a.m. and 8 p.m. On an average day, the maximum dispersal of *Pst* spores occurred between 1 p.m. and 3 p.m., coinciding with the highest average WV and global radiation, while the maximum dispersal of *Pt* spores occurred with a slightly earlier peak in the day, between 9 a.m. and 1 p.m.

**Figure 6.**
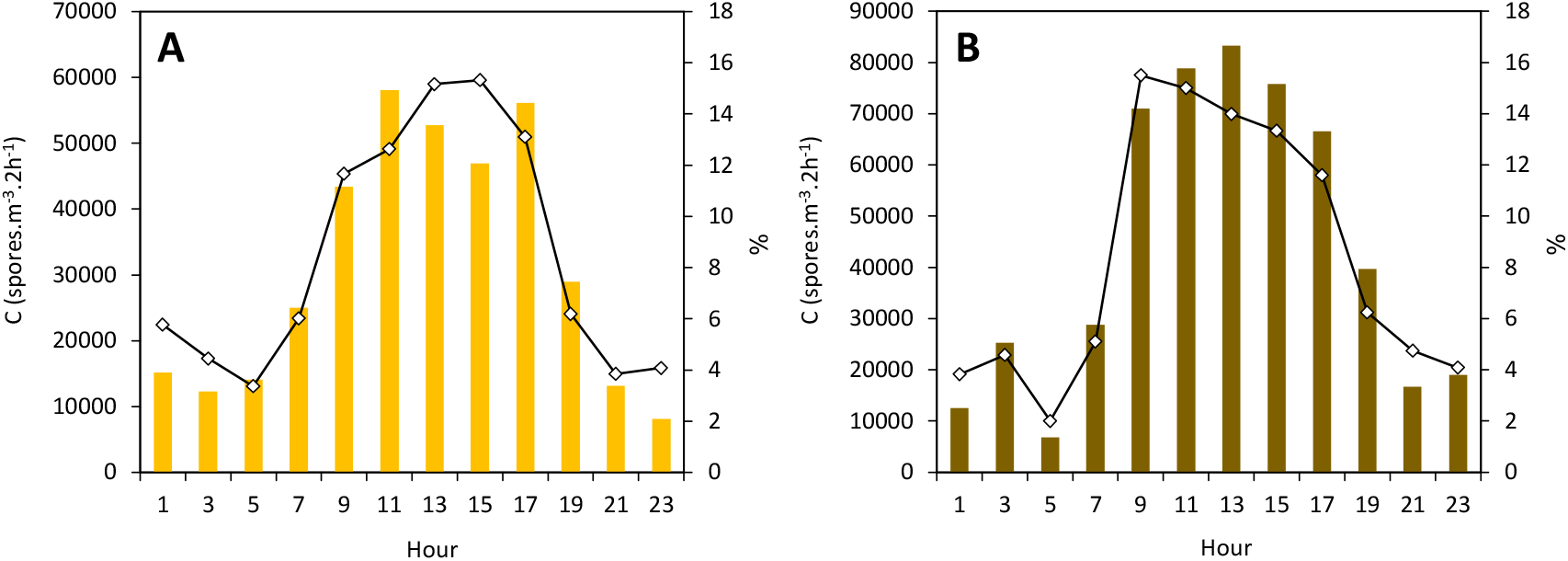
Intraday trend in airborne spore concentration of **(A)** *Puccinia striiformis* f. sp. *tritici* (*Pst*, stripe rust) and **(B)** *Puccinia triticina* (*Pt*, leaf rust) expressed in spores.m^-3^.2h^-1^ (bars) and in percentage of the total daily number of airborne spores trapped (lines) during the active epidemic phase. Average concentrations were estimated in 2-hour intervals (UTC hour) during the active epidemic phase. The active epidemic phase, which spans from DOY 125 to DOY 149 for *Pst* and from DOY 150 to DOY 186 for *Pt*, was defined as the interval from the first day when the total daily number of airborne spores exceeded 10000 to the last day when this quantity was trapped.

### General impact of meteorological variables on airborne spore concentration

The OLS model explained 13,8% of the variance for *Pt* and 7,9% for *Pst* (**Table 1**). A global F-test indicated that the explanatory variables had a statistically significant effect (for *Pt* F = 17.69, p < 0.0001; for *Pst* F = 10.25, p < 0.001). T had a positive impact for *Pt* (β = 231.811, p = 0.002) but not for *Pst* (β = -41.941, p = 0.575), RH had a negative impact for *Pst* (β = -78.480, p < 0.001) but not for *Pt* (β = 8.773, p = 0.672), and WV had a positive impact, significant for *Pt* (β = 476.567, p = 0.005) but not significant fo *Pst* (β = 237.319, p = 0.076). The effect of R, on the other hand, was positive and significant for both *Pt* and *Pst* (β = 1560.293, p < 0.001 and β = 690.110, p = 0.008, respectively). These results indicate that both WV and R, being systematically linked to increased airborne spore concentrations, are better predictors than RH and T, whose impact is more variable.

**Table 1.**
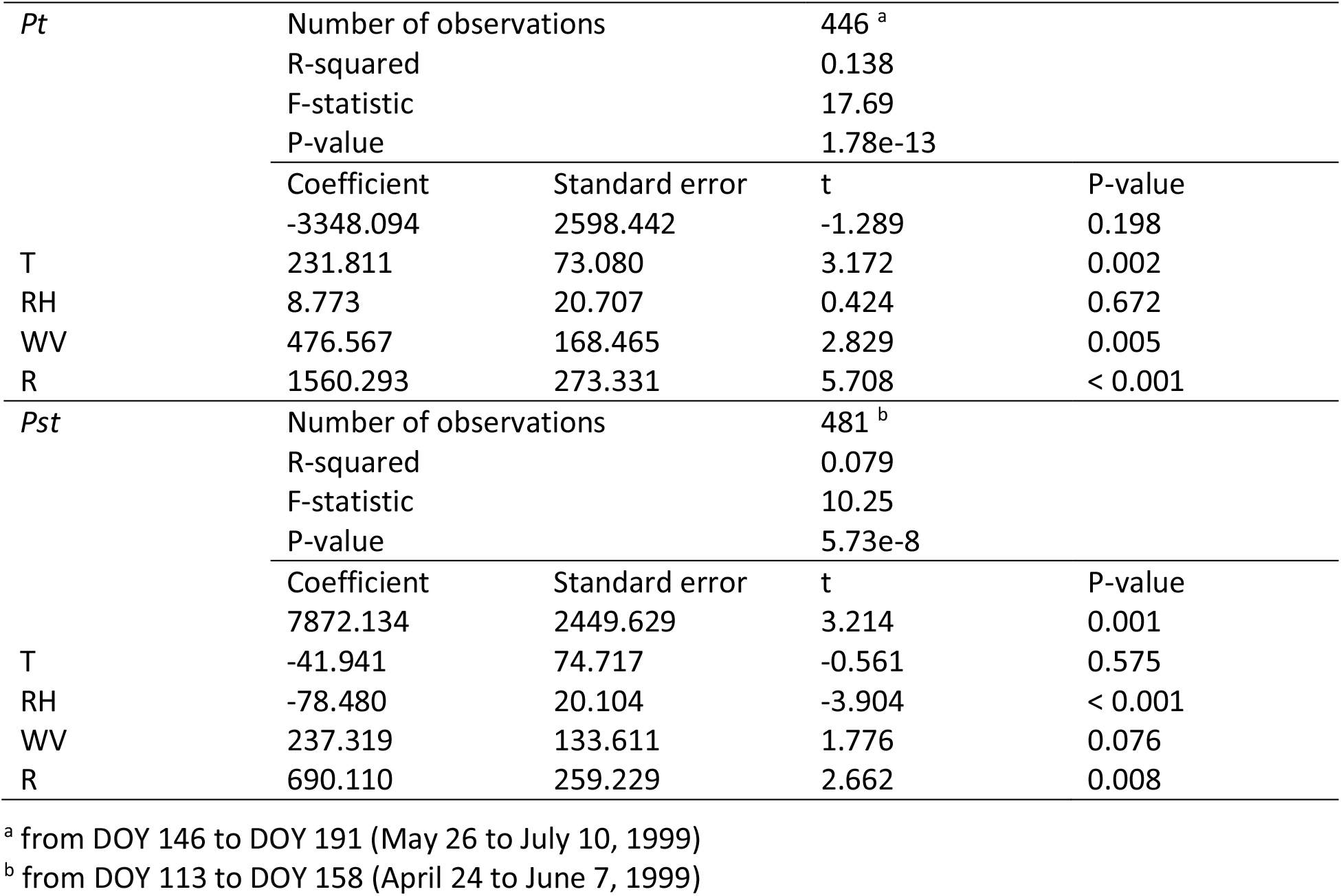
Impact of three climate variables (air temperature T, relative humidity RH, wind velocity WV, rainfall R) on 2-hour airborne spore concentration of *Puccinia triticina* (*Pt*) and *Puccinia striiformis* f. sp. *tritici* (*Pst*) based on an OLS regression model.

### Effect of rain events on the airborne spore concentration

The complete bi-hourly dataset presented in **Supplementary Figure S1**, setting airborne spore concentration against rain events, reveals the diversity of the rain impacts and allowed for the identification of two examples for each type of impact. Precursor rains almost consistently led to an increase in spore concentration. This increase was more pronounced when the rainfall was substantial (Type 1). Even rains of very low intensity could trigger a rise in spore concentration when they followed a prolonged dry interval (Type 2). Follower rains generally resulted in a decrease in spore concentration when they immediately followed one or more previous rain events (Type 3). When preceded by a 2 to 4-hour dry interval, a follower rain had more variable effects, sometimes leading to an increase in spore concentration (Type 4) and other times causing a decrease or a relative stability (Type 5), depending on the rainfall intensity. Sometimes during extended rain intervals, i.e. characterized by more than four successive 2-hour rain events, the diurnal periodicity of airborne spore concentration was heavily disrupted, with concentrations drastically reduced for over 24 hours (e.g., after DOY 155-156, DOY 158, and DOY 185-186 for *Pt*, and after DOY 167 and DOY 140 for *Pst*). Additionally, the last significant concentration peaks were consistently observed during a rain event (DOY 185 and DOY 187 for *Pt*, and DOY 147, DOY 149, and DOY 150 for *Pst*).

For *Pst*, 24 precursor rains and 33 follower rains were identified between DOY 127 and DOY 158 and included in the analysis. For *Pt*, 26 precursor rains and 35 follower rains were identified between DOY 147 and DOY 186. On average, the suspension effect from precursor rains was characterized by an increase in airborne spore concentration +2569% for *Pst* (**Figure 7A**) and +2436% for *Pt* (**Figure 8A**). A more limited effect was noted during follower rains, with a modest increase in spore concentration (+165% for both *Pt* and *Pst*), barely exceeding the ‘No-rain’ baseline (**Figures 7A** and **8A**). The analysis also revealed how spore concentration evolves during the interval following a 2-hour rain event. After precursor rains, spore concentrations showed few variations, with an additional increase of +81% for *Pst* (**Figure 7B**) and a decrease of -48% for *Pt* (**Figure 8B**), both averaging slightly different or even lower than the ‘No-rain’ baseline. After follower rains, changes in concentration were of a similar magnitude, with an additional increase of +29% for *Pst* (**Figure 7B**) and +48% for *Pt* (**Figure 8B**). These values represent average trends, encompassing both intervals where the rain had stopped and intervals where it continued, as the post-rain interval in practice also can be defined as a follower rain event. The classification of the rain effects into categories (< 0,]0-100], ]100-200], ]200-300], ]300-400], ]400-500], > 500%) confirmed the significance of rainfall characteristics (intensity, number of consecutive rain events, etc.). During more than half of the precursor rain events, the spore concentration of both rust species increased by more than 500%, and was followed by a decrease in almost three-quarters of the subsequent 2-hour intervals. During and after more than two-thirds of the follower rain events, the spore concentrations decreased.

**Figure 7.**
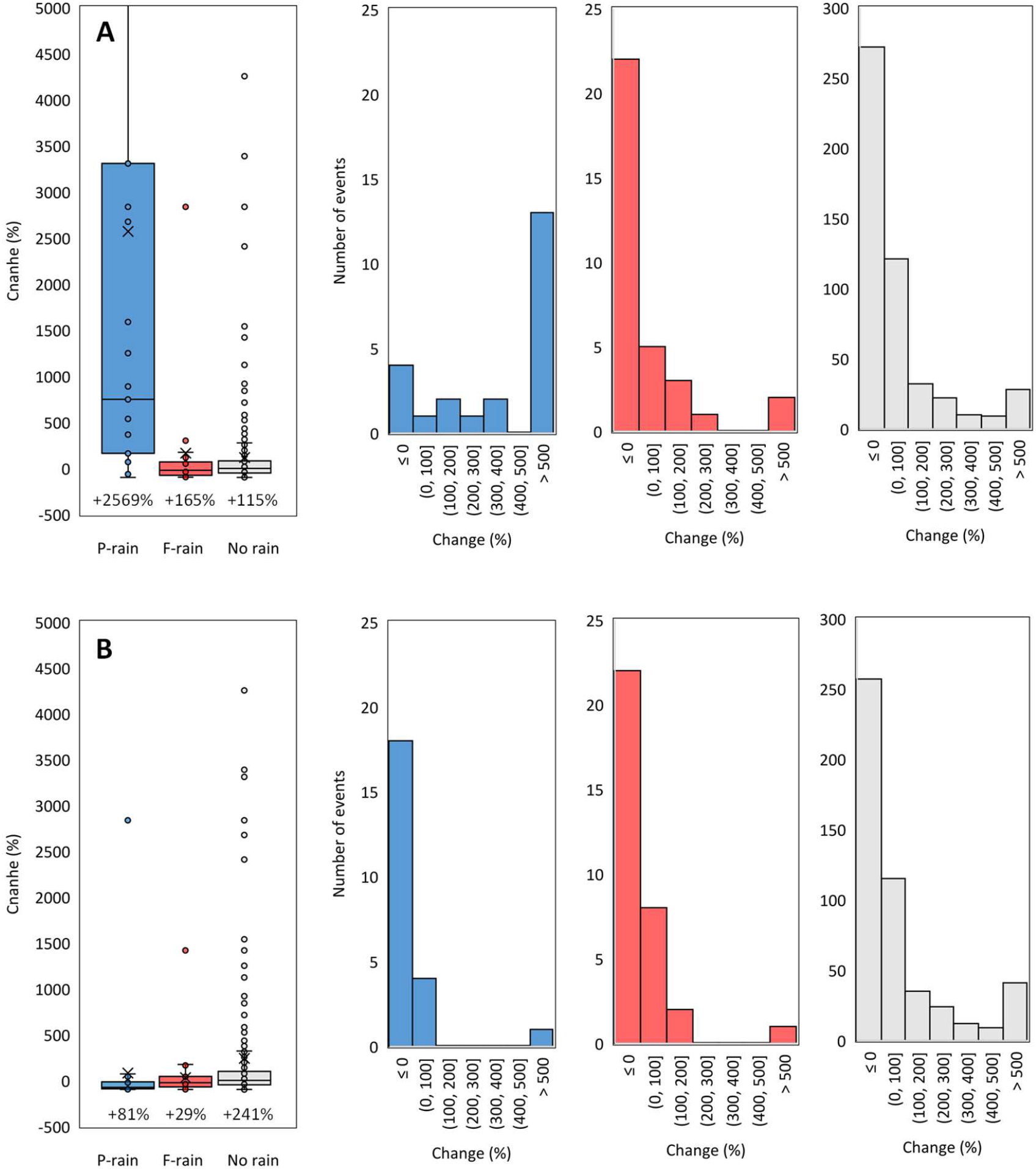
Impact of rain on airborne *Puccinia striiformis* f. sp. *tritici* spore concentrations, comparing changes induced by ‘precursor’ rain events (P-rain; blue bars) and ‘follower’ rain events (F-rain; red bars) to the average changes observed in the absence of rain (‘No rain’; grey bars). The measured impacts are distinguished as follows: **(A)** ‘suspension effect’ during the rain event, showing the percentage variation in spore concentration relative to the 2-hour interval preceding this rain event; **(B)** ‘wash-out effect’ just after the rain event, showing the percentage variation in spore concentration relative to the 2-hour interval following this rain event. The three rightmost graphs show the impact of all rain events, categorized by effect type.

**Figure 8.**
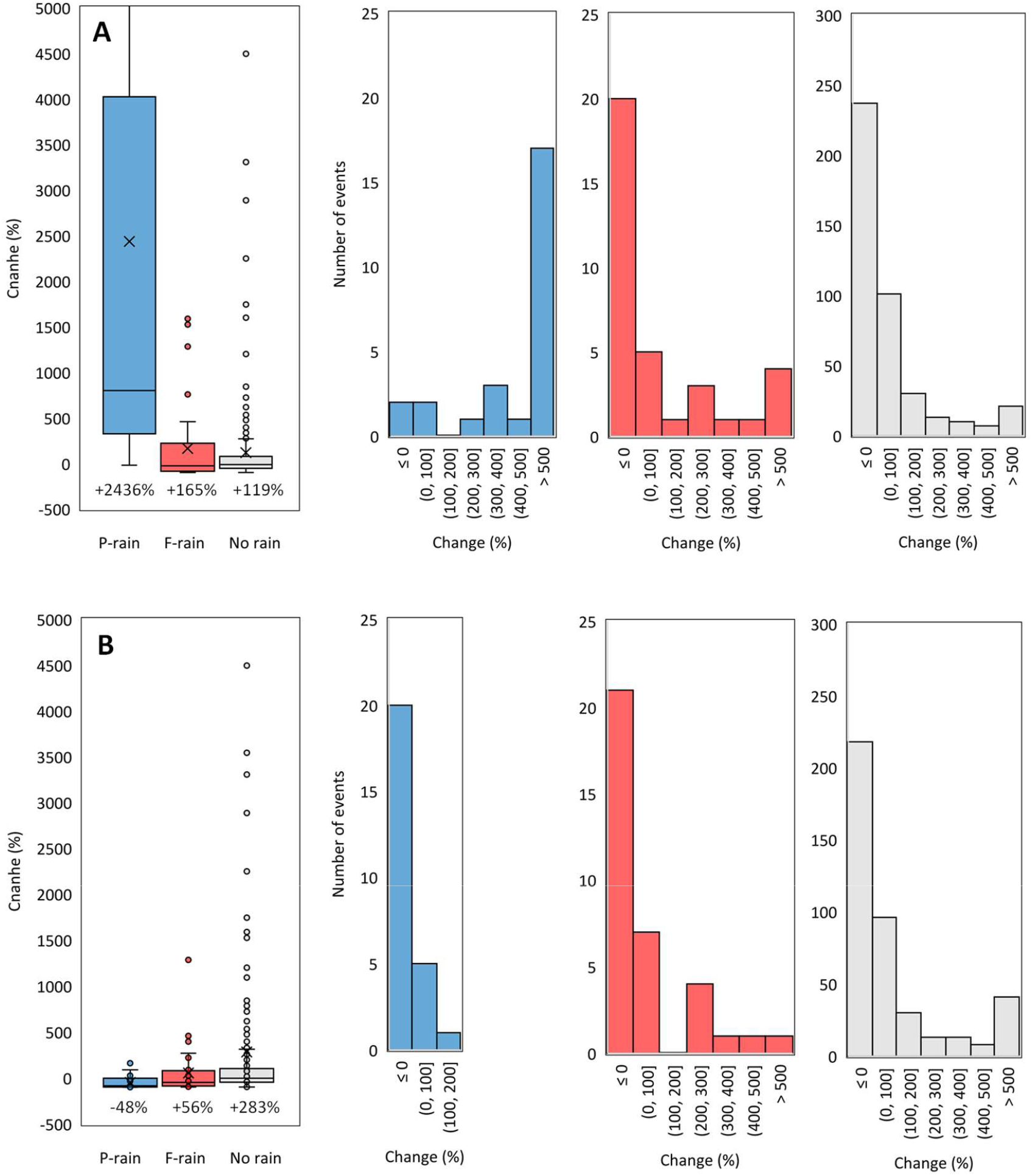
Impact of rain on airborne *Puccinia triticina* spore concentrations, comparing changes induced by ‘precursor’ rain events (P-rain; blue bars) and ‘follower’ rain events (F-rain; red bars) to the average changes observed in the absence of rain (‘No rain’; grey bars). The measured impacts are distinguished as follows: **(A)** during the rain event (‘suspension effect’), showing the percentage variation in spore concentration relative to the 2-hour interval preceding the rain event; **(B)** just after the rain event (‘wash-out effect’), showing the percentage variation in spore concentration relative to the 2-hour interval following the rain event. The three rightmost graphs show the impact of all rain events, categorized by effect type.

### Effect of humidity and wind on the airborne spore concentration

When categorized into value classes, gust intensity (GI) emerged as a relevant explanatory factor, despite exhibiting high variability. On average, spore concentration was ten times higher for gust levels greater than 0.35 compared to those below 0.15 (**Figure 9A**). The relationship with wind velocity (WV) was also increasing but reached a maximum. This threshold effect appeared slightly earlier for *Pt* (2.5-3 m.s^-1^) than for *Pst* (3-3.5 m.s^-1^) (**Figure 9B**). Regarding relative humidity (RH), the average airborne spore concentration peaked at levels between 50% and 65% for both rusts, with very low concentrations observed when RH exceeded 85% (**Figure 9C**). The high variability observed in the spore concentrations for each class of meteorological variable is linked to the diurnal periodicity of the removal process and should encourage caution when interpreting these results. When testing these relationships using meteorological variables averaged over the same 2-hour intervals of each day, a clearer relationship emerged (**Supplementary Figure 2**). However, these relationships inherently reflect the diurnal cycle of airborne spore concentration, higher during the day, as RH and WV exhibit similar but opposite fluctuations (lower HR during the day and higher WV at night; **Figure 10**).

**Figure 9.**
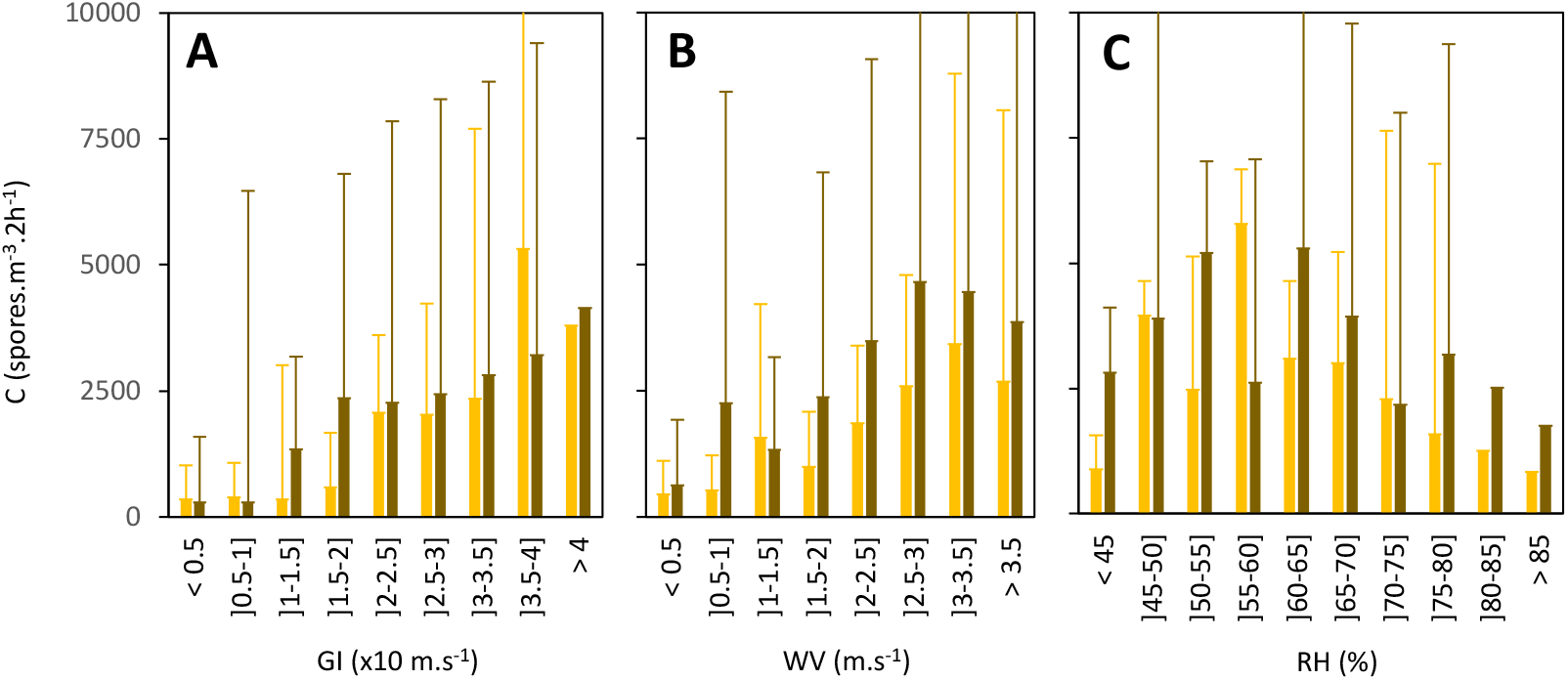
Effect of **(A)** relative humidity (RH, in %), **(B)** wind velocity (WV, m.s^-1^) and **(C)** guts intensity (GI, in x10 m.s^-1^) on the airborne spore concentration of *Puccinia striiformis* f. sp. *tritici* (*Pst*, stripe rust; yellow bars) and *Puccinia triticina* (*Pt*, leaf rust; brown bars) expressed in spores.m^-3^.2h^-1^ during the active epidemic phase, i.e. from DOY 125 to DOY 149 for *Pst* and from DOY 150 to DOY 186 for *Pt*. Error bars represent the standard deviation (SD) of the mean.

**Figure 10.**
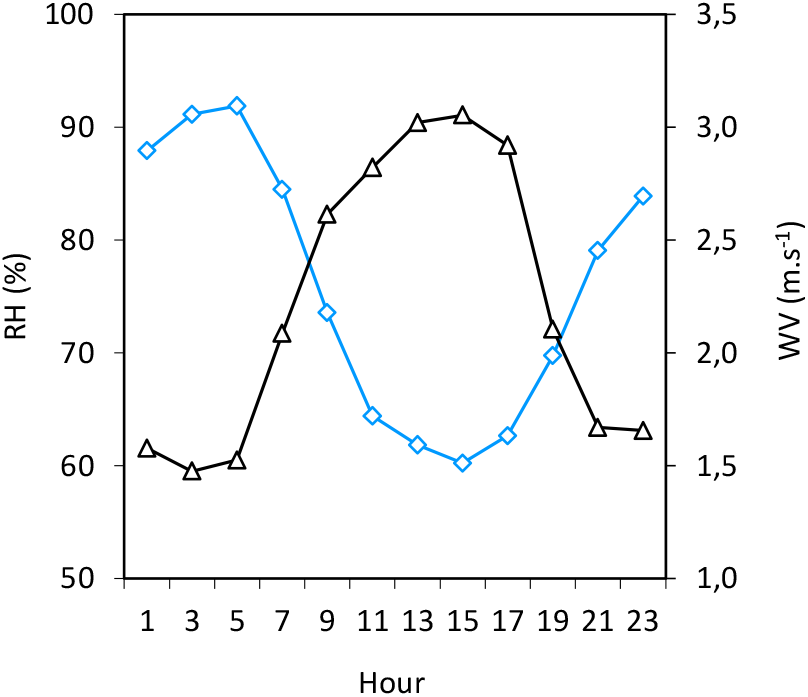
Intraday trend in relative humidity (RH in %, blue line) and wind velocity (WV in m.s^-1^, black line). Each point represents the average estimated in 2-hour intervals (UTC hour) from DOY 112 to DOY 186.

### Effect of humidity and wind velocity on the size of dispersal units

The size of *Pst* dissemination units that were trapped decreased as relative humidity increases: at 90% RH, over 70% of dissemination units consist of single spores, compared to less than 35% when RH was below 60% (**Figure 11A**). Conversely, the size of dissemination units increased with WV, with more than 80% being single spores when it is below 1 m.s^-1^, compared to less than 50% when WV exceeded 3 m.s^-1^ **(Figure 11C**). The effect of these two meteorological variables on the size of *Pt* dissemination units followed the same trend but was much less pronounced (**Figures 11B and 11D**).

**Figure 11.**
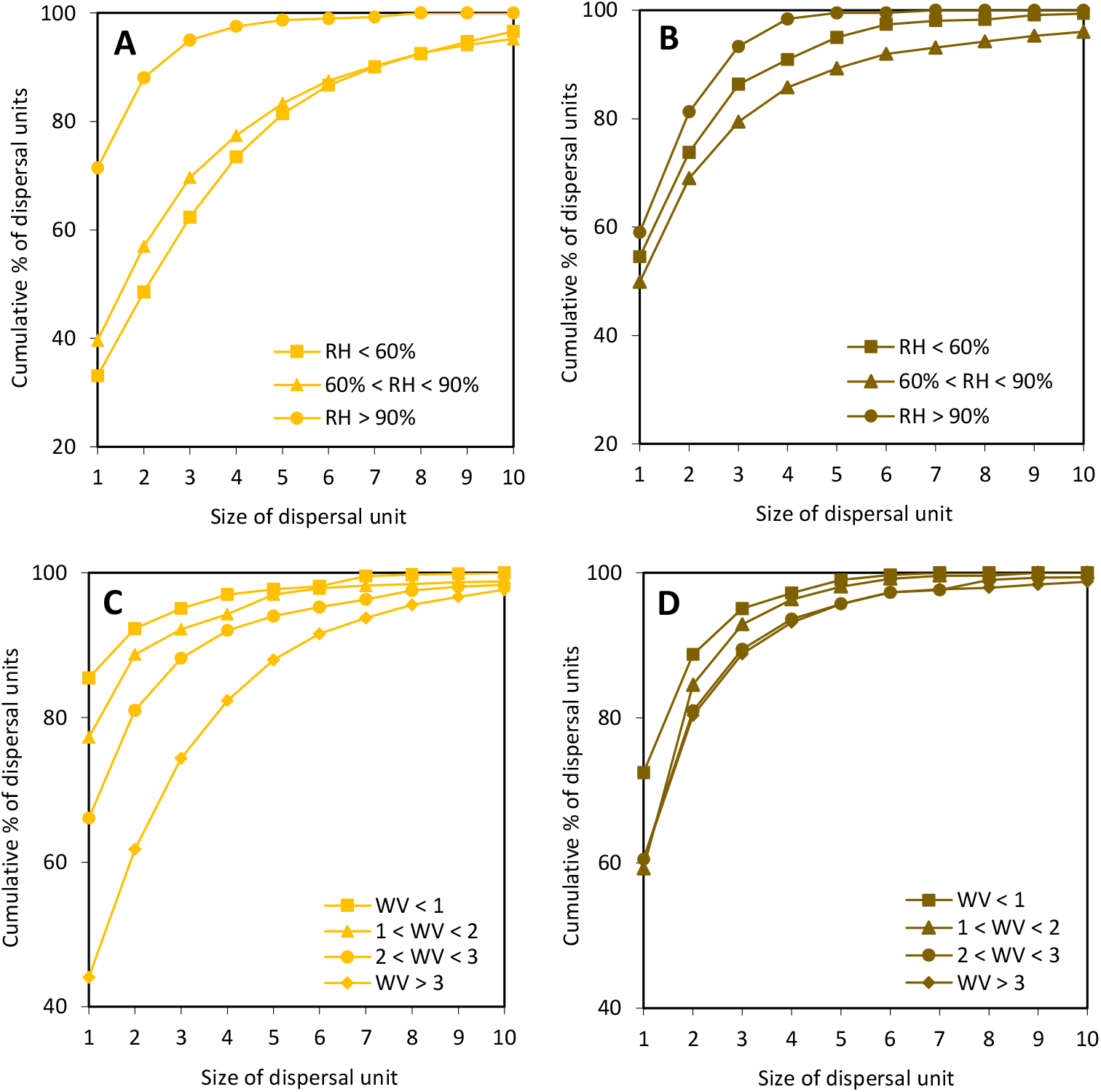
Influence of **(A, B)** relative humidity (RH, in %) and of **(C, D)** wind velocity (WV, in m.s^-1^) on the size of dispersal units (number of spores per dispersal unit) captured by the spore trap for *Puccinia striiformis* f. sp. *tritici* (*Pst*, stripe rust) and *Puccinia triticina* (*Pt*, leaf rust).

## Discussion

### Seasonal patterns and diurnal periodicity shape the general evolution of airborne spore concentrations

The leaf rust epidemic occurred approximately one month later than the stripe rust epidemic, consistent with the shift in the dynamics of airborne spore concentrations of *Pt* and *Pst*. This aligns with long-standing observations in France, where leaf rust typically appears later on upper leaves, usually between the emergence of the flag leaf and the heading stage, except in some southern areas. From a climatic perspective, rainfall types varied (intense but brief showers, prolonged rain, drizzle) and the total precipitation during from May 1 to July 1, 1999 (98.8 mm) matches the average over the past 30 years in the Yvelines (105.7 mm; MeteoFrance). Despite being based on a single year of data, experimental results can be considered representative of broader trends.

A two-tiered dynamic of airborne *Pt* and *Pst* spore concentrations emerged, characterized by: (1) a seasonal trend that aligns with the progression of the rust on wheat leaves, serving as a reservoir of spores whose quantities increase with the number of uredinia and then decrease as the uredinia become exhausted and the leaves senesce; (2) a daily trend characterized by a diurnal periodicity, correlated with fluctuations in climatic variables potentially influenced by sunlight, which reach their minimum or maximum around midday.

### Several meteorological variables influence the diurnal periodicity of airborne spore concentrations

To date, two main types of epidemiological studies have examined the impact of climate conditions on airborne spore concentrations. The first type focuses on correlations between spore concentrations and meteorological variables, usually with the goal of better understanding epidemic dynamics and improving forecasting models (e.g., Hjelmroos 1993; Gu et al., 2000; Pakpour et al., 2015; Sánchez Espinosa et al., 2024). The global regression (OLS), conducted here separately for *Pt* and *Pst*, falls within this category. It revealed a positive influence of wind and rain, and a negative influence of humidity, on the mechanisms driving spore removal. Temperature clearly emerged as having no strong effect at the time step considered, which aligns with my critique of the approaches proposed by Sánchez Espinosa et al. (2023) and Hu et al. (2024) in the Introduction. This clarifies why conclusions regarding the influence of rainfall (for instance, based on a “correlation between the density of urediniospores and total precipitation over 25 days before the spore peak”; Hu et al., 2024) can be meaningless, given the timescale of the rain-driven mechanisms, which occurs between minutes and hours rather than days or weeks. Only a second type of study that considers multiple physical mechanisms can adequately address the complexity of the effects of rain. Process-driven approaches require making assumptions and refining variables, such as distinguishing between precursor and follower rain events, and assessing their effects on spore concentration not only within the same bi-hourly period but also in the subsequent one. This approach yielded conclusive results in the current study.

The diurnal periodicity in airborne spore concentration in cereal rusts was established several decades ago (Hirst, 1953; Pady et al., 1965). This trait is common with many other pathogens such as *Uncinula necator* (Willocquet and Clerjeau, 1998) or *Cercospora arachidicola* (Alderman et al., 1987). Here, nearly 80% of spores were released during the day between 8 a.m. and 8 p.m., when relative humidity was lower, and average wind velocity was higher. The impact of these two factors, as drivers of daily periodicity of depletion and repletion of uredinia, was also demonstrated for *Puccinia arachidis* (Savary et al., 1985). The delay in the daily spore peak of *Pst* compared to *Pt* may be attributed to the differing influence of humidity on spore clustering in the two species.

### The impact of humidity and wind differs slightly between Pt and Pst likely due to the size of the dispersal units

The current results confirmed that relative humidity has a greater impact on *Pst* spore clustering compared to *Pt*. Under natural conditions, contrary to the findings of Rapilly and Foucault (1968) and a complementary experiment conducted under controlled conditions alongside this study (Suffert, 1999), the size of the trapped dispersal units for *Pst* decreased as relative humidity increased. This apparent contradiction may arise from the fact that during the day saturation deficits coincide with high wind velocities, which dry the atmosphere. The absence of wind thus limits the dispersal capacity and consequently the trapping of larger, heavier clusters of spores, even though they predominate. This hypothesis aligns with the effect of clustering on reducing spore dispersal distance (Ferrandino and Aylor, 1984; Chamecki et al., 2012) and the significant effect of wind on the amount of *Pst* spore trapped (**Table 1**), for which larger dispersal units were captured once wind speeds exceed 3 m.s^-1^ (**Figure 11C**), contrary to *Pt*. We should be aware of potential sampling bias introduced by the trapping process and the fact that the number and type of trapped propagules may not accurately reflect airborne spore concentrations.

The relationship presented in **Supplementary Figure S2**, while requiring cautious interpretation due to the data structure, allows for a clearer visualization of the differences between *Pt* and *Pst* compared to **Figures 9B** and **9C**. Specifically, it highlights a stronger positive influence of wind velocity and relative humidity on the airborne suspension of spores for *Pt*, for which the slope of the line passing through the minimum point is approximately 50% steeper than that of *Pst*. The heavier dispersal units of *Pst* have likely more difficulty being dispersed and trapped, even when relative humidity and wind velocity are more favorable. This aligns with the force required to suspend *Pst* spores, as measured by Geagea et al. (1997), and the presence of early foci and a steeper gradient tail for stripe rust compared to leaf rust (Chamecki et al., 2012; Rapilly, 1970).

### The effects of rain on the diurnal periodicity of airborne spore concentrations are disruptive, asynchronous, and antagonistic

Rain appeared as an asynchronous disruptive driver of dry dispersal. This impact depends on the characteristics of a rainfall (intensity, duration, and time elapsed since the previous one). The effects of precursor rains are consistently positive and easy to identify and quantify, with an average increase in airborne spore concentration by a factor of 25. In contrast, the effects of follower rains are more difficult to characterize because they result from antagonistic mechanisms.

The rain-driven mechanisms for dry spore dispersal (rain-puff) and wet spore dispersal (rain-splash, wash-out, wash-off) that explain fluctuations in airborne spores above a wheat canopy are summarized in **Figure 12**. Rain-puff occurs with greater intensity when uredinia are filled with spores, typically after periods of low wind and no rain for several days. Intense gusts further amplify this effect. Rainfall remove the available spores in the atmosphere and deposit them very quickly on leaves by wash-out. Prolonged rain events deplete spore source (uredina) and wash off spores already deposited on leaves. This representation aligns with the findings of Savary et al. (2004) regarding the effects of simulated rainfall in three tropical pathosystems: coffee rust, peanut rust, and bean angular leaf spot. The authors showed that rain-puff predominates in low-intensity, short rainfalls, while rain-splash remains stable in importance under high-intensity, long rainfalls. The data from the current field study confirm that the impacts of rain are antagonistic and depends on its intrinsic characteristics (intensity, duration) as well as the sequence of rain events (whether they occur close together or are spaced apart).

**Figure 12.**
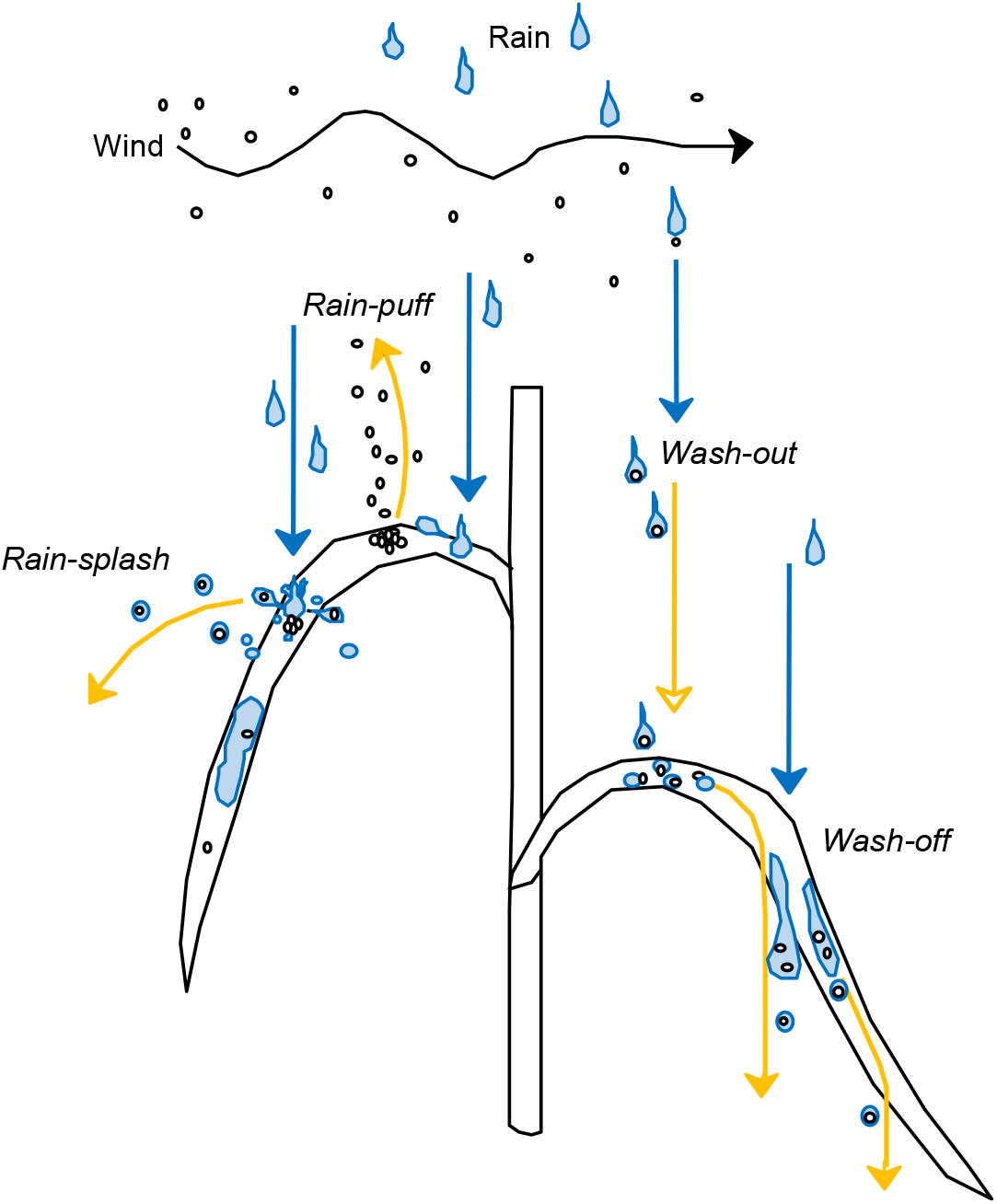
Four rain-driven mechanisms for dry spore dispersal (rain-puff) and wet spore dispersal (rain-splash, wash-out, wash-off) that explain fluctuations in airborne spores above a wheat canopy.

While rain-puff was clearly demonstrated by the current dataset, the wash-out was less evident, likely because this mechanism manifests later and more gradually. The fact that the increase in airborne spore concentration during the interval following a 2-hour rain event was of the same magnitude (or even lower) than the ‘No-rain’ baseline is indicative of its existence. The demonstration of the effects of wash-out and wash-off was conducted alongside the current field study through two analytical experiments: the first by estimating the concentration of waterborne spores collected in rainwater; the second by comparing rust severity on wheat seedlings exposed to rain with those not exposed (Suffert, 1999; Sache, 2000).

### Applications and potential improvements

Using airborne inoculum near-ground monitoring as a decision aid requires an understanding of the physics of particle transport within plant canopies (Mahaffee et al., 2023). The knowledge gained here can further contribute to forecasting models, as the quantified effects of wind and rain align well with what has been experimentally demonstrated through physical approaches. However, it also highlights the limitations of purely statistical, non-mechanistic analyses in capturing the nuanced effects of these variables on specific epidemiological processes. Especially, it underscores the limitations of correlative approaches that either use datasets with time intervals disconnected from the underlying processes, leading to the artificial detection of certain variables influences (e.g., Sánchez Espinosa et al., 2023; Hu et al., 2024), or face technical difficulties in obtaining trapping data at the same time resolution as meteorological data over extended periods (Chamecki et al., 2012).

Possible improvements relate to the precision of the meteorological dataset. The first potential improvement involves the measurement of gusts, which was not incorporated into the OLS model because this variable was derived from wind velocity and excluded due to inconsistencies in time steps with spore trapping. The second potential improvement concerns rainfall characterization, which was based here on the cumulative water height over 2-hour periods. This is more refined than in many previous studies but does not allow for a detailed physical characterization of the rainfall. To obtain information on the size and velocity of the falling droplets, as well as the distribution of their kinetic energy—crucial for modeling dispersion trajectories through splash—an optical spectro-pluviometer can be utilized (Salles et al., 1999; Saint-Jean, 2003). Such precision is only beneficial if spore quantification can be finely synchronized, meaning it must occur at time intervals much shorter than one or two hours, as was done in the current study. This requires improvements in trapping technology and spore counting methods. If trapping bands are retained, image analysis tools are available for the automatic quantification of spores (Lei et al., 2018), although distinguishing between species remains a challenge. For this reason, molecular diagnostic tools represent a significant avenue for improvement. This is especially relevant given the considerable progress made since 1999, particularly with qPCR (Dedeurwaerder et al., 2011; Duvivier et al.; Araujo et al., 2020; Gu et al., 2020; Rodríguez-Moreno et al., 2020), as well as metabarcoding performed without prior assumptions (e.g., Chen et al., 2024). However, these technical advances should not overshadow the importance of considering the right timing and duration of spore trapping.

## Conclusion

Paradoxically, the more numerous the mechanisms driving an epidemiological process and examined under controlled conditions, the more challenging it becomes for an experimenter to investigate them holistically under field conditions, particularly when these mechanisms have antagonistic effects. However, investigating them in the field, in all their complexity, is crucial. Until now, spore dispersal— from removal to deposition—has been quantified individually under controlled conditions, without revealing which mechanisms dominate under specific rain conditions or what type of rain exerts which effects in field settings. The dataset analyzed in this study addressed this gap. Despite significant technological advances in spore quantification, the temporal pace for trapping remains a limiting factor if not carefully considered, which can hinder progress in plant disease epidemiology. Basing analysis on mechanism-driven hypotheses and avoiding inconsistent indiscriminate correlations between variables should be central to the research strategy.

## Acknowledgments

This work was a part of my agronomy engineering internship, which I defended on September 29, 1999, in Paris. Narrowly saving the archive box containing this 25-year-old dataset from the landfill during the move of the INRAE BIOGER laboratory from Grignon to Palaiseau in September 2022 made me realize the value of such experimental data. Inspired by this near-loss, I curated and reanalysed the data with fresh perspectives, more experience, and a touch of nostalgic clarity. I would like to thank all the colleagues who contributed to the data acquisition, particularly Ivan Sache and Laurent Huber for guiding me during my internship, as well as Sébastien Saint-Jean, Alain Fortineau, Brigitte Durand, Henriette Goyeau, Claude de Vallavieille-Pope, and Marc Leconte, for their technical support in Physics, Micrometeorology, and Plant Pathology.

## Supplementary Material

**Supplementary Figure 1.**
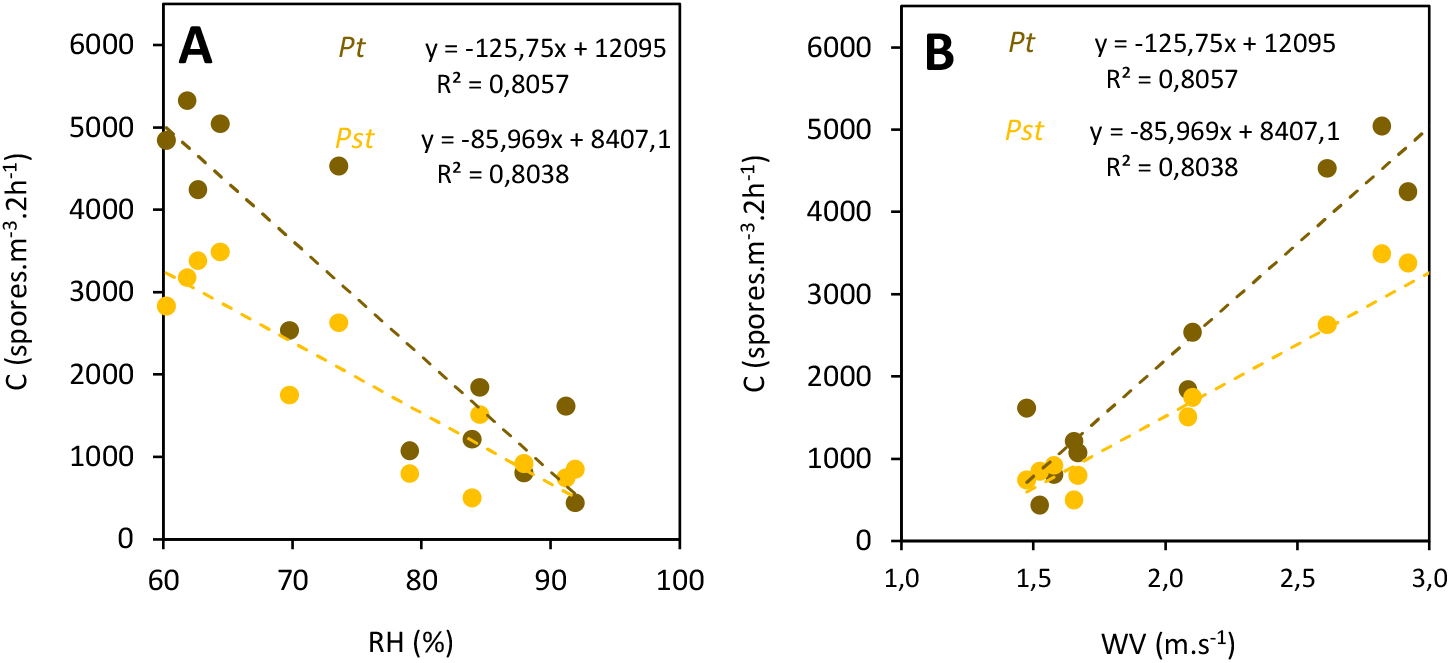
[Suppl. Fig S1 Suffert 2024.xls]. Airborne spore concentration C (spores.m-3.2h^-1^) of **(A)** *Puccinia triticina* (*Pt*, brown rust; brown bars) from days 146 to 191 (May 26 to July 10, 1999) and **(B)** *Puccinia striiformis* f. sp. *tritici* (*Pst*, stripe rust; orange bars) from days 113 to 158 (April 24 to June 7, 1999), in parallel with rainfall (mm.2h^-1^) and wind velocity (m.s^-1^).

**Supplementary Figure S2**. Relationship between the average airborne spore concentration of *Puccinia striiformis* f. sp. *tritici* (*Pst*, stripe rust) or *Puccinia triticina* (*Pt*, leaf rust) expressed in spores.m^-3^.2h^-1^ and **(A)** the relative humidity (RH, in %) or **(B)** the wind velocity (WV, in m.s^-1^). Each point represents the average of the variables estimated in the same 2-hour intervals of each day in the active epidemic phase, i.e. from DOY 125 to DOY 149 for *Pst* and from DOY 150 to DOY 186 for *Pt*.

## Data Availability Statement

Data are available in the INRAE Dataverse online data repository at https://doi.org/10.57745/WADAIY

